# Innate face detectors in the nidopallium of young domestic chicks

**DOI:** 10.1101/2024.02.15.580445

**Authors:** Dmitry Kobylkov, Orsola Rosa-Salva, Mirko Zanon, Giorgio Vallortigara

**Affiliations:** Centre for Mind/Brain Science, CIMeC, University of Trento, Rovereto, Italy

**Author notes:** Corresponding author: Giorgio Vallortigara P.zza Manifattura, 1, 38068 Rovereto, Italy; **Email:**.

## Abstract

Soon after birth, naïve animals and newborn babies show spontaneous attraction towards faces and face-like stimuli with three dark features representing eyes and a mouth/beak. While neurons selectively responding to faces have been found in the inferotemporal cortex of adult primates, face-selective domains in the brains of young monkeys seem to develop only later in life after exposure to faces. This has fueled a debate on the role of experience in the development of face-detector mechanisms, since face preferences are well documented in naïve animals, such as domestic chicks reared without exposure to faces. Here we demonstrate that neurons in a cortex-homologue area of one-week-old face-naïve domestic chicks selectively respond to a face-like configuration. Our single-cell recordings show that these face detectors do not respond to alternative configurations or isolated facial features. Moreover, the population activity of face-selective neurons accurately encoded the face-like stimulus as a unique category. Thus, our findings show that face detectors are present in the brains of very young animals without pre-existing experience.

## Introduction

Face perception is a fundamental set of cognitive skills^1^. In fact, our visual system is so primed to faces that we can recognize face-like patterns even in inanimate objects like clouds or the Moon, a phenomenon known as pareidolia. This is true not only for humans: for other animals too faces and facial features are highly salient, playing an important role in social interactions (mammals^2^, birds^3–5^, fish^6^, and even insects^7^).

Over the last four decades, significant progress has been made in understanding the neural mechanism of face perception. Neurons specifically responding to monkey and human faces have been found in the inferotemporal cortex (IT) of adult primates^8–9^ and the fusiform gyrus of humans (for a review see^10^). These neurons show a strong selectivity to faces compared to other visual or auditory stimuli. A crucial question, however, is whether this selectivity is only the result of experience or whether there are evolutionarily predisposed innate mechanisms for face detection.

Several lines of evidence support the idea that face detection mechanisms might be present from birth. Multiple studies have shown that behavioral responses to faces emerge very early in development. Newborn babies are attracted by a schematic upright face-like pattern with two eyes symmetrically placed above the mouth^11–12^. This behavioral bias was observed already in the uterus, where human fetuses reacted more to the upright face than to the inverted stimulus^13^. In a precocial species, domestic chicks, a similar preference for face-like stimuli has been found soon after hatching before they ever saw a face^4^. At the neural level too, face-selective neural responses have been recorded in the electroencephalogram (EEG) of newborn babies^14^ and the hemodynamic response (fMRI) of young infants^15^.

However, in contrast, fMRI evidence from monkeys suggests that face-selective domains (inferotemporal areas that are specialized for face processing) develop through extensive early-life experience with faces. Face domains were absent/underdeveloped in monkeys younger than three months^16^ or raised without exposure to faces^17^. These results have fueled a debate on the origins of face-selective neural responses, which is far from being resolved.

For instance, the hypothesis of an acquired mechanism of face detection does not account for the evidence of behavioral biases towards faces. Spontaneous face-preferences emerge before the development of the face-selective cortical domains^17^ and do not require pre-exposure to faces^4–5^. On the other hand, the face-naïve monkeys tested by Arcaro and colleagues^17^ were deprived of visual experience with faces for several months. Such long-lasting deprivation from biologically relevant sensory stimuli is known to alter cortical development^18^. The underdevelopment of the face domains in face-naïve monkeys could simply reflect this prolonged deprivation, rather than prove the absence of an evolutionary predisposed, innate neural mechanism tuned to faces. Thus, exposure to faces during sensitive developmental periods might be necessary for the maintenance rather than for the development of face detectors. In addition, the fMRI itself is probably not sensitive enough to detect single-cell face-selective responses in very young animals, which may precede the formation of face-domains. For example, in infant monkeys, Rodman et al.^19^ did find face neurons, albeit with firing rates substantially lower than in adults.

Thus, in the current discussion on the developmental origin of face detection, one of the main stumbling points is that so far none of the studies could detect single-cell neural responses to faces in newborn face-naive animals^20–21^.

It is important to stress that most of the studies that investigated neural responses to faces employed realistic face images. However, studies in human newborns and newly hatched chicks show that the innate response to faces is triggered mainly by the geometrical configuration of facial features, which may be considered as a “supernormal stimulus” *sensu* ethology^22^.

By using newly hatched chicks, we were able for the first time to study innate neural mechanisms for face detection at the level of single-unit responses, while avoiding long deprivation phases. Moreover, for our study we employed a schematic face-like stimulus, featuring two dark spots symmetrically positioned above the mouth/beak, forming an upside-down triangle within the head silhouette (Fig. 1B). This configuration triggers robust behavioral preferences and is recognized as ’face-like’ from early development through adulthood^23^. This simplified stimulus also controls for the influence of high-level visual features and allows assessment of the impact of individual facial features and their configurations.

**Fig. 1.**
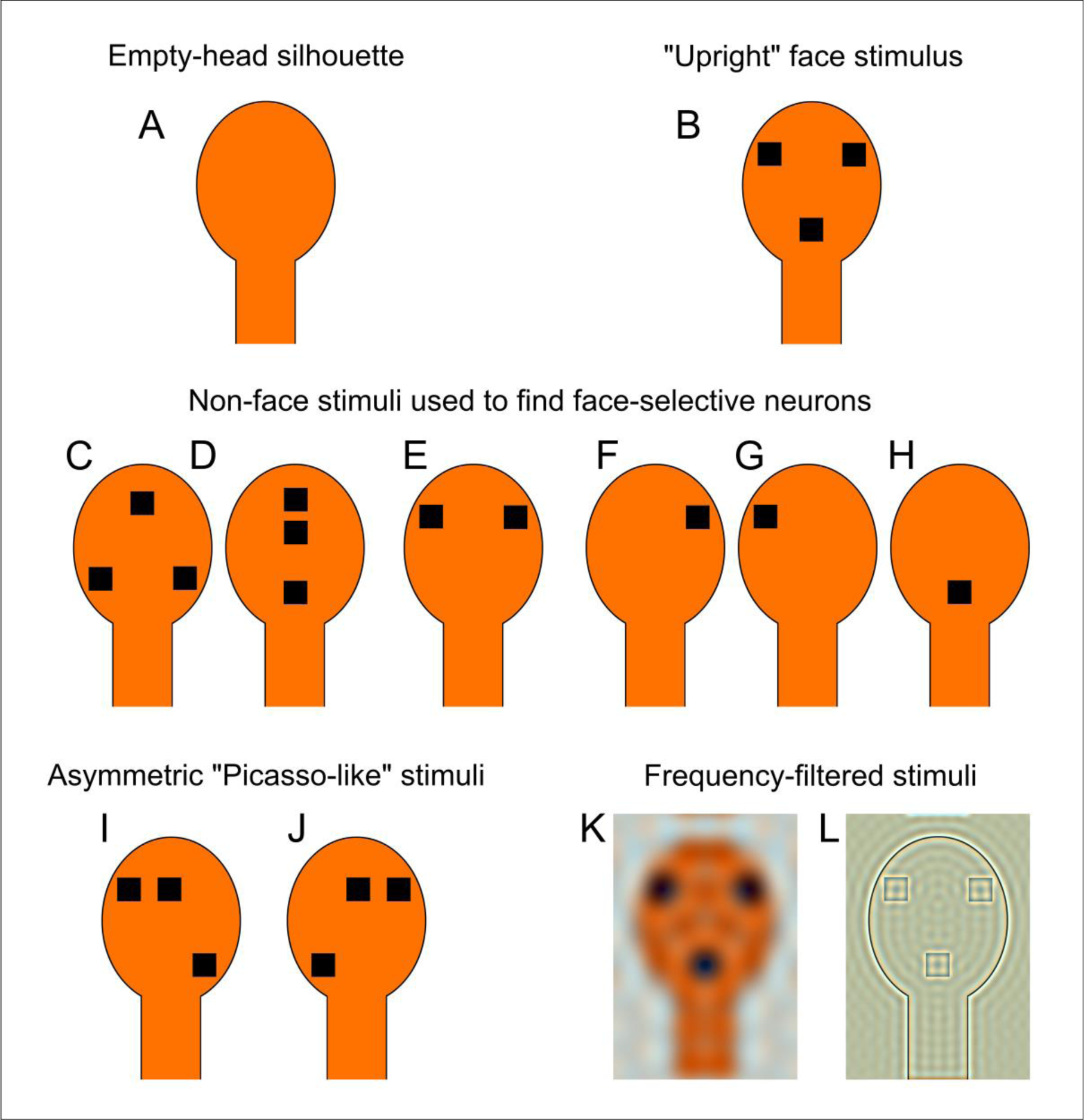
Experimental stimuli. (A) An empty head silhouette used for habituation and as a background image during stimuli presentation. Thus, every trial only the configuration of dots inside this head region (B-H, K-L) or a frequency component (I-J) of the stimulus was changing. We defined a face detector based on its significantly stronger response to the “upright” face-like configuration (B) than to other geometrical configurations with three dots (“inverted” (C) and “linear” (D)) or to single facial features: “eyes” (E), “left eye” (F), “right eye” (G), “beak” (H). Furthermore, we tested the response of putative face neurons to asymmetrical “Picasso-like” stimuli with eyes shifted from the medial sagittal line to the right (“picassoR” (I)) or to the left (“picassoL” (J)). In addition, we tested how face neurons respond to face-like stimuli dominated by low-frequency (K) or high-frequency (L) components.

With this approach, here we demonstrate that neural face detectors can emerge without specific prior experience by recording electrophysiological responses to face-like stimuli in young chicks that had never been exposed to faces. To identify face-selective neurons in young chicks, we compared single-cell electrophysiological responses to several configurations of face-like stimuli. Putative face-selective neurons responded more strongly to the “upright” stimulus with the normal configuration of facial features compared to either an incomplete face (“eyes”, “beak”, “left eye”, “right eye”; Fig. 1 E-H) or a different configuration of facial features (“inverted”, “linear”; see Fig. 1 C-D). We further characterized the response of identified face-selective neurons to asymmetric stimuli (“Picasso-like” faces), which violate the symmetrical configuration of the face template (Fig. 1I-J). Based on existing behavioral evidence, we expected innate face responses to be reduced for these asymmetric stimuli in chicks^4^ (but see^24^). Furthermore, we tested whether face-selective neurons would be sensitive to the spatial frequency content of the face-like stimulus, as innate face detectors have been hypothesized to selectively respond to the low-spatial frequency content of the image^25^. Consequently, we expected face-selective responses to be reduced for high-pass filtered images (“HF”) compared to low-pass (“LF”) filtered ones (Fig. 1K-L).

## Results

### 8% of NCL neurons selectively respond to the “upright” face configuration

We recorded neural activity from 540 neurons in the caudolateral nidopallium (NCL) of young face-naïve domestic chicks. 8% of all recorded neurons (N = 43) responded significantly stronger to the “upright” face-like stimulus, than to single facial features (“eyes”, “beak”, “left eye”, “right eye”) or to a different arrangement of these features (“inverted”, “linear”, see Fig. 2A and Movie S1 showing exemplary face-selective neural responses). The amount of these face-selective neurons was significantly higher than the chance level calculated based on the analysis of shuffled trials (3.7% of false-positive units; proportion test: p < 0.001, χ2 = 12.09, Fig. 2B). While the majority of stimulus-selective neurons preferred the “upright” face configuration, we also observed selective responses to other tested stimuli. Selective responses to the “beak” (7.2%, N = 39) occurred significantly more often than expected by the chance level (proportion test: p = 0.003, χ2 = 8.54). The number of selective responses to facial features in other configurations (“inverted” and “linear”, 6% N=33 each) was just above the chance level (proportion test: p = 0.04, χ2 = 4.16).

**Fig. 2.**
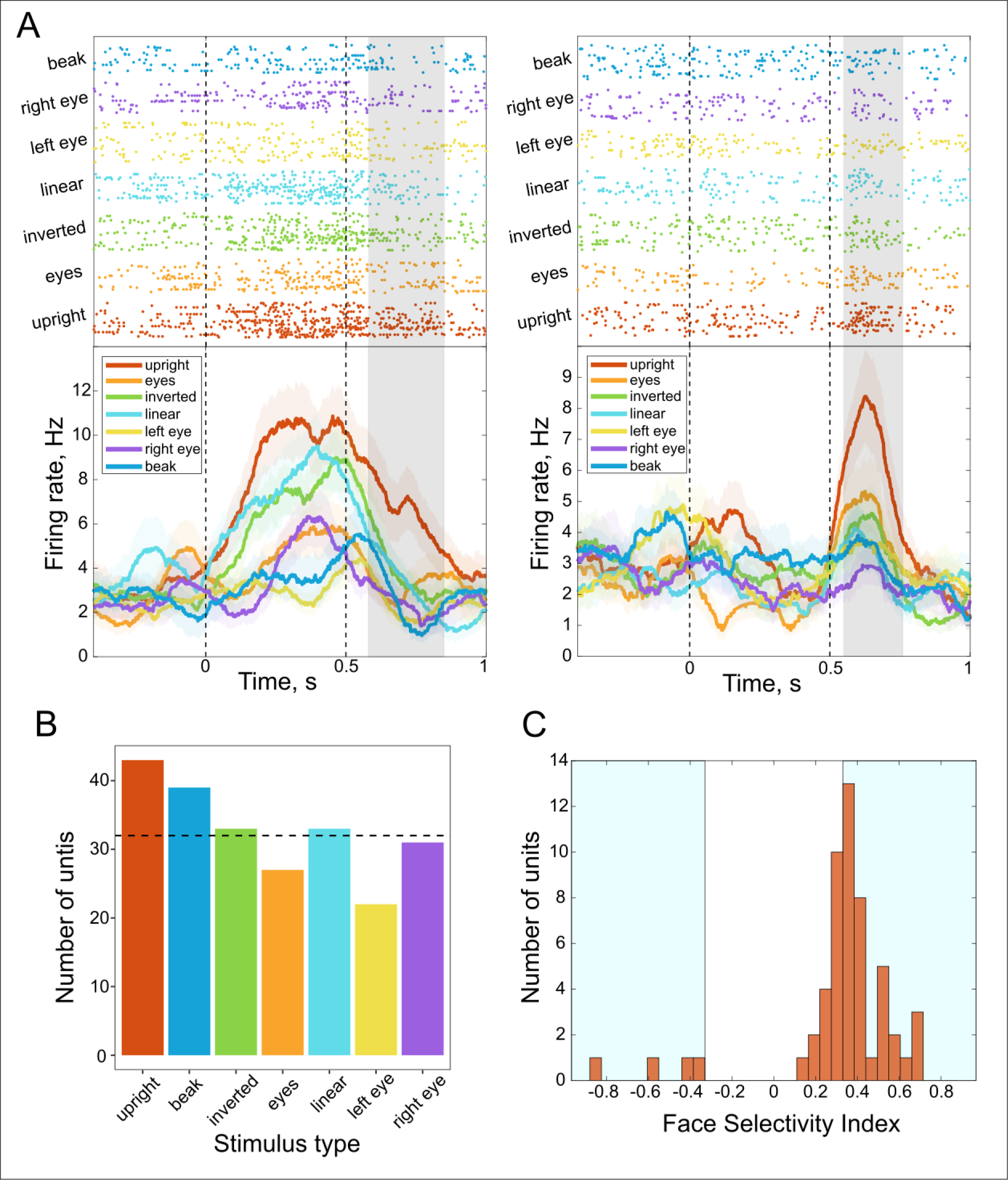
Face-selective neurons in the NCL of young domestic chicks. (A) Two example units that responded significantly stronger to the “upright” stimulus. Raster plots in the upper part show trial-by-trial neural responses grouped by stimuli types (shown in different colors). Lines of the raster plot represent single trials; dots correspond to neural spikes. The peristimulus time-histograms (PSTH) in the lower part represent the average neural responses to face-like stimuli smoothed by the Gaussian kernel (100ms sigma). The shadowing in the PSTHs corresponds to the standard error of the mean neural response. The grey shadow indicates the face-selective time window. (B) Number of units that showed selective responses to different configurations of face-like stimuli. The dashed-line shows the significance threshold calculated based on the analysis of shuffled trials (alpha = 0.05). (C) Face-selectivity index (FSI) quantifies the selectivity of individual face-responsive cells. A FSI >= 0.33 or <= -0.33 (shown with light-blue shadows) corresponds to a 2:1 ratio between face vs. non-face responses and is considered a strong selectivity.

During the face-selective response window, face-responsive neurons showed a strong preference towards the “upright” face configuration compared to other stimuli (the face-selectivity index (FSI) = 0.41 ± 0.14, Mean ± Standard Deviation). The majority of them (77%, N = 33) had the FSI of 0.33 or higher (Fig. 2C), which means that their response to the “upright” face was at least two times stronger than the averaged response to other non-face stimuli. Only 9% (N = 4) of face-responsive neurons showed negative FSI, which corresponds to them being selectively inhibited by the face stimulus.

The face detectors in the NCL of chicks were sensitive to the symmetry of the face stimulus (LME p < 0.001 (*numDF* = 5, *F* = 26.29) for the interaction factor “stimulus type” * “FSI”, Fig. 3). Units with the positive FSI, which increase their firing rate in response to the “upright” stimulus, show significantly decreased response to the “Picasso-like” images, both when the eyes were shifted leftwards and rightwards from the medial sagittal line (post-hoc Tukey test: p < 0.001 for “upright” vs. “picassoL” (*z* = 9.27) and “upright” vs. “picassoR” (*z* = 9.56)). Conversely, units with the negative FSI show significantly less inhibition in response to the “Picasso-like” stimuli (post-hoc Tukey test: p = 0.014 (*z* = -3.13) for “upright” vs. “picassoL” and p = 0.046 (*z* = -2.73) for “upright” vs. “picassoR”).

**Fig. 3.**
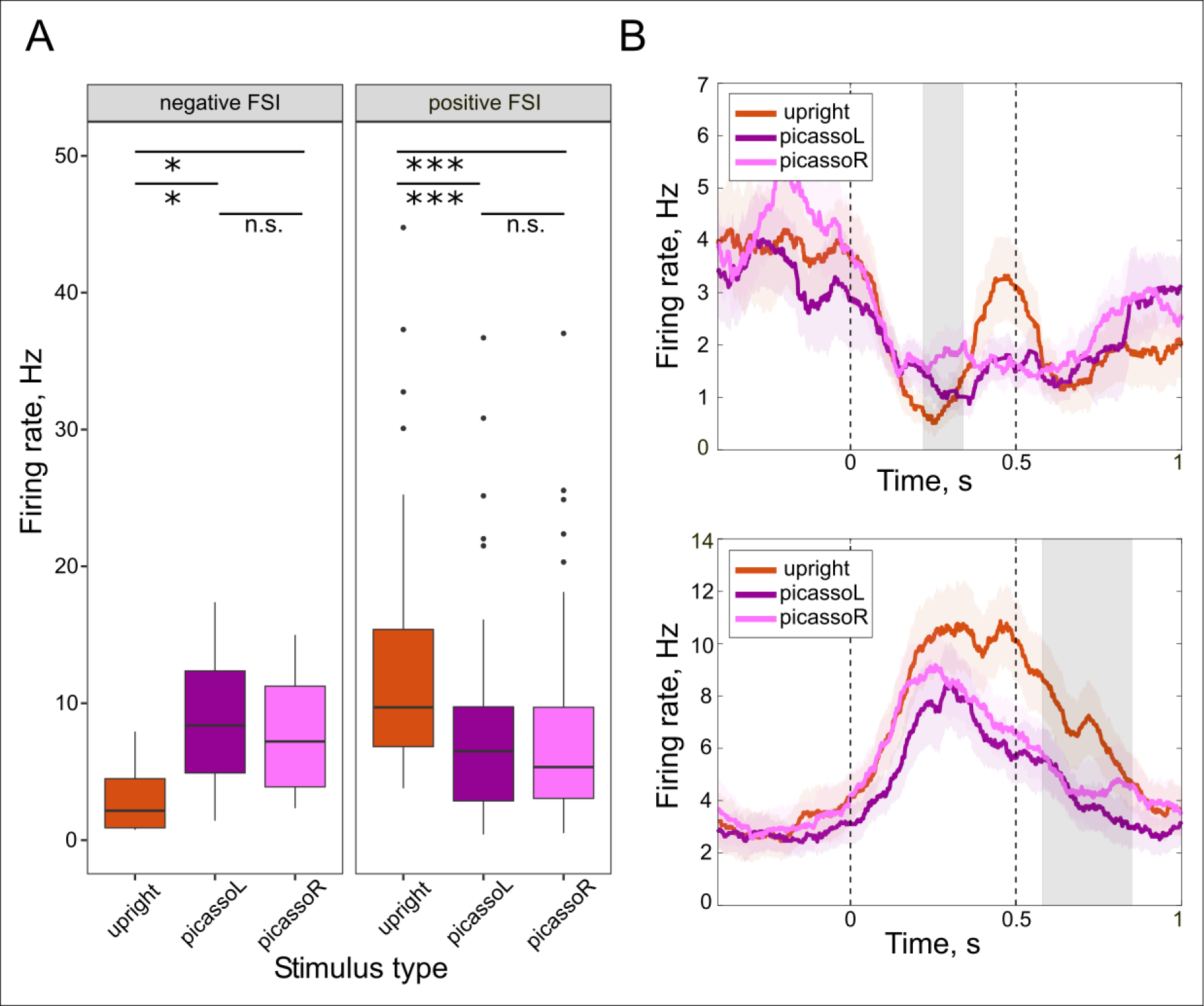
The face-selective units showed sensitivity to the symmetry of the face stimulus. (A) The average firing rate within the face-selective window was significantly different between the “upright” symmetric face and “Picasso-like” stimuli with eyes shifted either to the left (“picassoL”) or to the right (“picassoR”). Moreover, this effect was opposite in units with the negative face-selectivity index (FSI) and positive FSI. Face-selective cells with the negative FSI showed reduced inhibition, while cells with the positive FSI showed reduced excitation to “Picasso-like” stimuli. Significance levels are based on the post-hoc Tukey test of the Linear Mixed Effect model: * p < 0.05, *** p < 0.001, n.s. – not significant. (B) PSTHs of two exemplary neurons with the negative FSI (above) and the positive FSI (below), which show different response to the asymmetric stimuli. For the description of the PSTHs see Fig. 2.

We additionally compared the neural responses of the face-selective cells to the images with the low-and high-frequency components. The response strength was significantly different between LF and HF stimuli (LME, p = 0.02 (*numDF* = 3, *F* = 3.3) for the interaction factor “stimulus type” * “FSI”, Fig. 4). Units with the negative FSI show significantly more inhibition in response to the “LF” than to “HF” stimulus (post-hoc: p = 0.032 (*z* = -2.6)). However, contrary to our expectations, there was no significant difference between these stimuli in the response strength of FSI positive units (post-hoc: p = 0.24 (*z* = 1.75)).

**Fig. 4.**
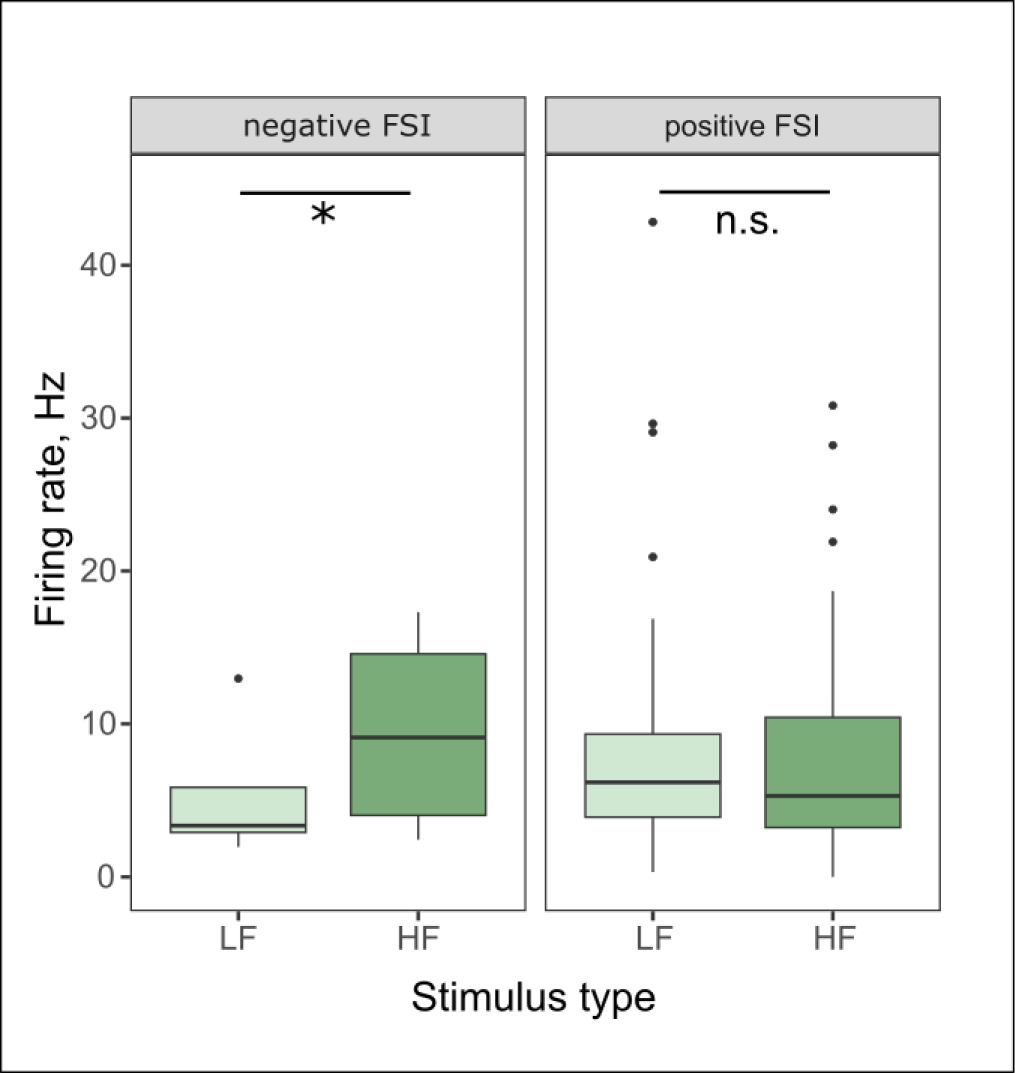
Neural responses of the face-selective cells to stimuli with the low-frequency (LF) and high-frequency (HF) components. The average firing rate within the face-selective window was significantly different between the “LF” and the “HF” only in units with the negative FSI. Significance levels are based on the post-hoc analysis of the Linear Mixed Effect model: * p < 0.05, n.s. – not significant.

### Population analysis

The face-selective window of face neurons was distributed relatively evenly throughout the time window of the stimulus presentation. Many neurons had a long response latency, starting even after the stimulus offset (shown in the heatmap in the Fig. 5). To evaluate the amount of information about the face-like stimuli contained in the entire population response over the period of the trial, we calculated the percentage of the variance explained (PEV) by the factor “stimulus type”. Stimulus-specific information content had two significant windows: it first increased significantly between 130ms and 380ms, and had another peak between 430ms and 630ms after the stimulus onset (Fig. 5). It means that the stimulus-specific information remained in the population response even after the offset of the stimulus.

**Fig. 5.**
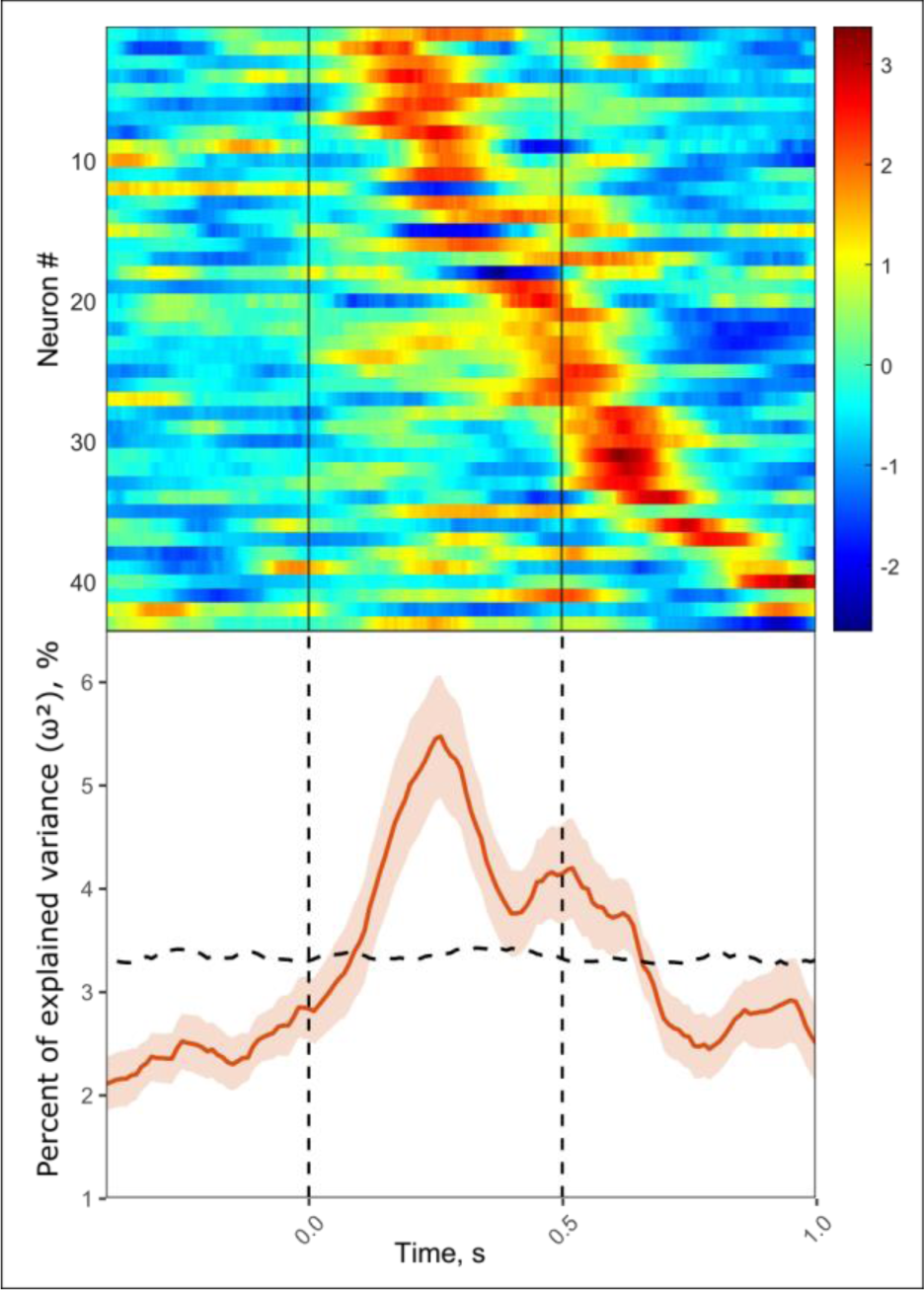
Population analysis of face-selective neurons. The heatmap in the upper part shows the neural response of all face-selective neurons (N = 43): each line represents the neural activity of a single neuron. The raw neural activity was first smoothed by the Gaussian kernel (100ms sigma), then averaged over all trials when the “upright” face stimulus was presented, and z-scored. The color map ranges from deep blue to dark red, which corresponds to low and high firing rate, respectively. The lower part of the figure shows the percentage of the variance explained (ω²) by the factor “stimulus type” during the whole period of the trial (see Methods). The horizontal dashed line shows the 95th percentile of the ω² from shuffled data (corresponds to p < 0.05). Vertical lines delineate the onset and the offset of the stimulus.

### Decoding analyses

We trained support-vector machines (SVMs) on the neural responses of face-selective units to estimate whether activity of this population can support an accurate categorization of face-like stimuli. The SVMs trained on the face-selective response windows were able to successfully categorize the “upright” face with high accuracy (42%, proportion test: p<0.001 (χ2 = 581.09), compared to the chance probability of 9%, Fig. 6A). The only stimuli that were mistakenly categorized as faces were asymmetric “Picasso-like” images (15%, p < 0.001, both for “picassoL” (χ2 = 35.05) and “picassoR” (χ2 = 32.03)). Additionally, the same model was able to categorize “HF” and “LF” stimuli, which seemed to form a distinct category. “Inverted” and “Picasso-like” stimuli were also uniquely coded, meaning that the number of correct estimations for these categories exceeded the number of false assignments.

**Fig. 6.**
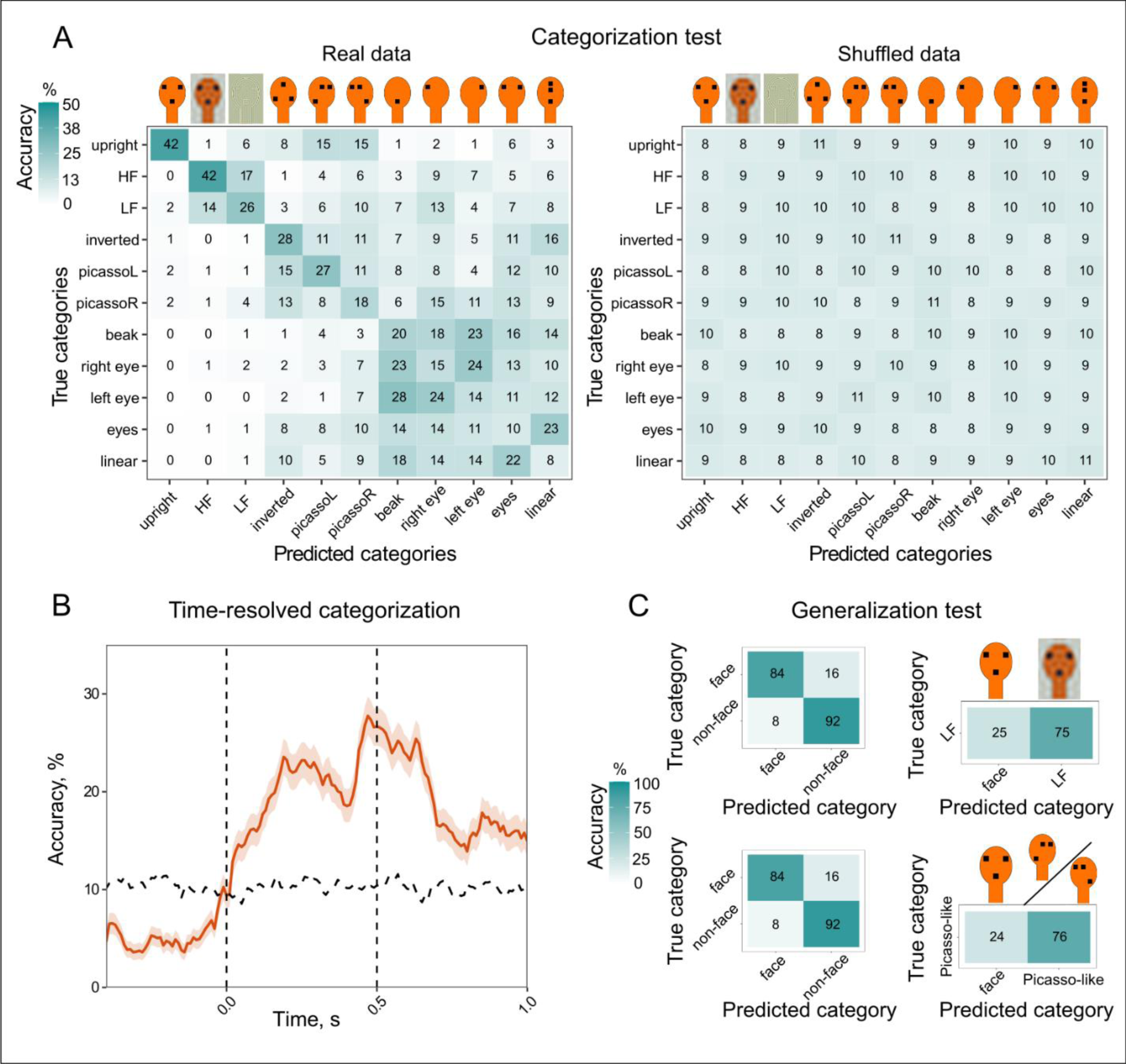
Decoding analyses based on supervised training of support-vector machines (SVMs). (A) The multi-class classifier was trained on the trial-by-trial average firing rate within face-selective response windows. Confusion matrix (on the left) shows the decoding accuracy of the classifier (in %). When the classifier was trained on shuffled data, its accuracy dropped to a chance level (confusion matrix on the right). (B) The time-resolved analysis of decoding accuracy of the “upright” face-stimulus. When the classifier was trained on the firing rate of consecutive 10ms time bins, its accuracy increased significantly above the chance level already 30ms after the stimulus onset. The chance level (horizontal dashed line) was estimated as 95% confidence interval calculated based on shuffled data. The vertical dashed lines delineate the onset and the offset of the stimulus. (C) Generalization test shows performance of the binary classifier trained on all-but-one stimuli types. First, classifier was trained to categorize stimuli into “face” and “non-face” classes based on the average firing rate within the face-selective window (upper and lower left confusion matrix). Then, the classifier was asked to categorize the new stimulus type (e.g., “LF”: upper right confusion matrix; “Picasso-like” stimuli: lower right confusion matrix).

To estimate how the accuracy of decoding changes over the time of the stimulus presentation, we trained SVMs on the consecutive 10ms time bins for the whole duration of the trial. Interestingly, the accuracy of face-decoding was significantly above the chance level starting already from 30ms after the stimulus onset (Fig. 6B). Moreover, the accuracy remained significantly high during the whole period of the trial, even after the offset of the stimulus.

SVMs trained on all stimuli types showed high decoding accuracy for “upright” face vs. other stimuli. Hence, we performed an additional generalization test by training SVMs on a subset of stimuli types to perform a binary face vs. non-face classification (Fig. 6C). Subsequently, we tested the performance of the SVMs on the stimuli types not included into the training to see whether these remaining stimuli would be spontaneously classified as faces or non-faces. We performed the generalization test consecutively on the following stimuli: “eyes”, “inverted”, “linear”, three-dots-stimuli (“inverted” and “linear” together), one-dot-stimuli (“left eye”, “right eye”, “beak”), Picasso-like stimuli (“picassoL” and “picassoR” together), “HF”, and “LF”. All tested stimuli were classified with high accuracy as non-faces (“eyes”: 93%; “inverted”: 83%; “linear”: 95%; three-dots-stimuli: 89%; one-dot-stimuli: 96%; Picasso-like stimuli: 76%; “HF”: 89%; “LF”: 75%). However, it is interesting to note that “Picasso-like” and “LF” stimuli were classified as faces in ca. 25% of all cases, which is significantly higher than the proportion of false-assignments during the test of the classifier (proportion test: p < 0.001, “Picasso-like” stimuli: χ2 = 173.92; “LF”: χ2 = 194.5).

## Discussion

Single-cell responses to faces have not been previously described in newborn face-naïve animals. Here we show that in one-week-old domestic chicks never exposed to faces, ca. 8% of neurons in the NCL selectively respond to a schematic face stimulus. These neurons constitute a neural population that encodes face-like stimuli as a separate category, as demonstrated by the analyses of the population response (PEV) and decoder performance.

While the idea of innate face detectors in the brain has been debated for a long time, the experimental evidence in support of this theory remained rather scarce (reviewed by^26^). Face-specific neural responses have been detected in the EEG of newborn babies already during the first days after birth^14^. Still, even at this age in human babies, it is extremely difficult if not impossible to exclude the influence of exposure to faces. On the other hand, studies on face-deprived monkeys^2, 17^ might be prone to artefacts related to possible side effects of long-lasting face deprivation.

By using precocial domestic chicks, we were able to minimize the face-deprivation period to a few days only and record neural activity in very young face-naïve animals. The observed face-selective neural responses correspond well with the spontaneous behavioral bias towards faces known in newly hatched chicks^4^. This provides strong evidence that the face detection mechanism in chicks is an inborn feature of the brain.

Contrary to the hypothesis of innate face detectors, Arcaro and Livingstone^27^ suggested an alternative explanation: early-life selectivity for faces could be attributed to an innate hierarchical proto-organization of the visual system. This innate hierarchical map in monkeys would show selective responses only to low-level features like shape and spatial frequency. This organization might serve as a scaffold for further development of category-selective responses. This hypothesis, however, does not seem to explain our results. Our face-selective neurons showed significantly stronger responses to the “upright” face compared to the “inverted” and the “linear” configurations. Importantly, the “upright” face stimulus is virtually identical to the “inverted” and the “linear” ones in terms of spatial frequency and luminosity (fig. S1). Therefore, the selectivity of the neural response can originate solely from the specific face-like geometrical configuration. It is possible, however, that this innate and potentially evolutionary conserved mechanism might be not directly involved in the recognition of individual conspecifics. Instead, mechanisms for spontaneous detection of face-like patterns likely serve to drive animals’ attention to social stimuli, e.g., to facilitate imprinting in newborn animals. Therefore, the functional role of the face-selective neurons for individual recognition in chicks has yet to be elucidated.

Notably, so far, all attempts to find face cells in the avian brain have yielded underwhelming results^28–30^. However, all previous studies in adult birds have used naturalistic images of faces. These images of birds/human faces are visually complex, with many local features, making it difficult to control relevant properties of the stimuli such as spatial frequency and luminosity. At the same time, even naturalistic face images still lack many aspects that might be important for a putative face detector: these are static images, presented on a flat screen, with colors that are unnatural to birds’ vision. Since we do not know the relative importance of all these features for birds, they might perceive such “naturalistic” images as rather unnatural and confusing.

Contrary to realistic face images, the schematic face-like pattern employed in our study served as a “supernormal” stimulus eliciting a strong selective response in the innate face detectors (Fig. 1B). The face detectors were sensitive not only to the particular number of visual elements (two eyes and a beak), but also to the specific geometric configuration and orientation of these elements. Their response to the geometrically identical, but inverted face was significantly reduced compared to the “upright” face stimulus with eyes placed above the beak. Similarly, in newborn human babies, inversion of the face stimulus also resulted in a weaker EEG response^14^. The face inversion effect observed at the neural level underlies a well-known behavioral phenomenon: inverted faces were shown to attract less attention in human babies^12^ and in young chicks^4^.

The difference in the neural response between “upright” and “inverted” faces cannot be explained by an attentional bias towards top-heavy stimuli, where more visual elements are concentrated in the upper part of the image^24, 31^. In behavioral experiments, newly-hatched chicks spontaneously prefer the normal-face configuration over the equally top-heavy “linear” stimulus^4^. Likewise, in our study, neurons selective for the “upright” face configuration responded significantly less to the “linear” top-heavy stimulus. This confirms that the neurons we describe in the NCL of chicks are selective to the distinct face-like configuration, rather than to single visual elements or their arrangement *per se*.

Apart from the upright triangular configuration, the vertical symmetry of facial features is one of the main properties that might facilitate face detection. Face symmetry has been utilized to enhance algorithms for automatic face detection *in silico*^32^. The face detectors that we recorded in the brains of chicks also relied on face symmetry. The face-selective neurons responded weaker to “Picasso-like” stimuli with eyes shifted to the side of the medial sagittal line. Accordingly, in behavioral experiments with face-naïve chicks, no preferences were found between asymmetric faces and control stimuli^4^. Interestingly, the analysis of the SVM classifier performance revealed similarities between “Picasso-like” and symmetric “upright” stimuli at the level of neural population response. The spontaneous categorization of “Picasso-like” stimuli as something between a face and a non-face object has also been reported in humans. In newborn babies, the EEG response to asymmetric stimuli was closer to the normal “upright” face than to the “inverted” configuration^14^. In human adults, fMRI revealed slightly different responses to Picasso’s paintings with realistic and asymmetric faces^33^.

Several studies have suggested that low-to mid-spatial-frequency components play a pivotal role in fast face detection^34–35^ (reviewed by^36^). In our recordings, units with a negative FSI (inhibited by “upright” faces), similarly reduced their response to the “LF” compared to the “HF” stimuli. Furthermore, “LF” stimuli were treated as faces with a relatively high probability of 25% by a classifier trained to categorize faces vs. non-faces. Hence, the neural population response to normal and low-frequency filtered faces shared some similarities. In the units exhibiting a positive FSI, however, we did not observe significant differences between the neural responses to “LF” and “HF” stimuli. The overall small difference between “LF” and “HF” stimuli might be explained by the reduced contrast they present compared to other stimuli due to filtering and subsequent luminance adjustment. Decreased contrast might have affected perception of the frequency-filtered stimuli. This is indirectly supported by the analysis of the classifier performance, which shows a high level of similarity between “LF” and “HF” stimuli. Hence, the frequency-filtered stimuli, which differ in their contrast from the rest of the stimuli, could have triggered a specific neural response.

It is important to stress that the relative role of various spatial frequency components in face detection remains largely unexplored, particularly at the single-cell level. Rolls et al.^37^ observed that face neurons in the superior temporal sulcus of monkeys respond to both low-pass and high-pass filtered faces, although with reduced amplitude. Results from EEG^38^ and MEG^35^ also suggest that fast and accurate face detection depends on the combination of both high and low-spatial frequency information. Moreover, according to fMRI data, the low-spatial-frequency component of faces might be primarily processed already at the subcortical level^39^. We, on the other hand, performed our extracellular recordings in the NCL, which is a higher-order processing area of the avian pallium functionally analogous to the mammalian prefrontal cortex^40^. Hence, we might expect that the differences between “LF” and “HF” stimuli would be more pronounced at the earlier subcortical stages of processing^25, 41–42^.

The hypothesis that detection of faces might occur already subcortically has not been directly tested at the level of single-cell responses in face-naïve newborn animals. However, responses to schematic face-like stimuli have been found in the superior colliculus of adult monkeys^41–42^. Our analyses of the SVM performance also offer indirect evidence that holistic detection of faces may occur in the subcortex. First of all, in the time-resolved analysis, the classifier successfully decoded the “upright” face stimulus shortly after the stimulus onset, which suggests the rapid detection of the face by the visual system. Second, in the categorization test, the “upright” face configuration did not overlap with responses to single facial elements. If the face-selective response would require separate detection and integration of single facial features (eyes and a beak), one could expect these stimuli to elicit a neural response similar to the “upright” face stimulus in the population of face-selective neurons. This was not the case, which indicates holistic processing of the face-like configuration. However, direct evidence for an innate subcortical face-processing mechanism can be provided only by neural recordings from subcortical areas in face-naïve young animals.

In addition to face-selective responses, our study revealed a considerable number of neurons responding to stimuli of different types. The number of selective responses to “linear” and “inverted” configurations was just above the chance level (p = 0.04), so it might be premature to make any assumptions about their potential biological function. However, a much larger percentage of all recorded neurons (7%) showed a strong selectivity for the “beak”. The fact that these neurons did not respond to other one-element stimuli (“left eye” and “right eye”) suggests that their neural response was primarily triggered by the distinct position of the beak within the head silhouette. For birds, the beak is an important facial feature that provides a variety of signals, e.g., during feeding behavior (gulls^43^) or sexual displays (pigeons^3^). Also in fowls, tidbitting serves as a salient cue for the presence of food, as a part of mating display in adults and of a parental display performed by brooding hens^44–45^. Whether the observed beak-selective neural responses might be related to an innate component of these behaviors in domestic chickens remains to be established.

In summary, our study represents a significant step in revealing the innate neural mechanism of face detection. Neurons in the NCL of domestic chicks entirely devoid of any early-life experience with faces showed a strong selective response to the face-like stimuli. This means that the face detectors we observed in the chicks’ brains are innate and do not require any pre-existing experience.

## Methods

### Subjects

For the study 6 domestic chicks (*Gallus gallus domesticus*) from the Aviagen ROSS 308 strain were used. Fertilized eggs from a local commercial hatchery (CRESCENTI Società Agricola S.r.l.–AllevamentoTrepola–cod. Allevamento 127BS105/2) were incubated and hatched within incubators (Marans P140TU-P210TU) at 37.7 °C and 60% humidity in a dark room. After hatching in dark incubators, chicks were isolated and housed individually in metal cages (28 cm wide × 32 cm high × 40 cm deep) with food and water available ad libitum, at a constant room temperature of 30–32 °C and a constant light–dark regime of 14 h light and 10 h dark. All experimental protocols were approved by the research ethics committee of the University of Trento and by the Italian Ministry of Health (permit number 539/2023-PR).

### Experimental setup

Experiments were performed in a rectangular shaped arena (34 X 54 X 27 cm; W X L X H) with wooden walls. One wall of the arena was replaced by a computer screen (AOC AGON AG271QG4, 144Hz) used for stimuli presentation. The arena was divided in two sections by a metal grid placed 31 cm from the screen preventing chicks from directly approaching it. A custom-build automatic reward system consisted of a feeder with mealworms, whose lid was attached to a servo motor and controlled by Arduino Uno. Stimulus presentation and reward was controlled by Bonsai software with BonVision toolbox^46–47^.

### Habituation procedure

To prevent chicks from seeing any faces, all habituation and experimental procedures were performed while wearing a full-face mask painted black. On the 2nd day post hatching, chicks learned to peck on mealworms. Between the 3rd and the 6th day after hatching the chicks were habituated to the setup. First, chicks were habituated to an empty head region (Fig. 1A) appearing on the screen. Initially, the birds received mealworms (the lid of the feeder opened for 500 – 1000ms) every time a stimulus appeared on the screen, which motivated them to pay attention to any new image appearing on the screen. During subsequent habituation, we gradually decreased the reward probability down to 30-40%, so that birds would still pay attention to the screen even without getting a mealworm. This procedure allowed us to minimize rewarding during actual recording sessions.

### Surgery and recordings

On the 7th day after hatching chicks were fully anaesthetized using Isoflurane inhalation (1.5 – 2.0% gas volume, Vetflurane, 1000mg/g, Virbac, Italy) and placed in the stereotaxic apparatus with a bar fixed at the beaks’ base and tilted 45° to ear bars. Local anaesthesia (Emla cream, 2.5% lidocaine + 2.5% prilocaine, AstraZeneka, S.p.A.) was applied to the ears and skull skin before and after the surgery. Metal screws were placed into the skull for grounding and stabilisation of the implant. A small craniotomy was made in the skull above the NCL (1.0 mm anterior to the bregma, 4.5 mm lateral to the midline, Fig. S2) on the right hemisphere (5 chicks) or on the left hemisphere (1 chick). For extracellular recordings we used self-wired tetrodes made out of formvar-insulated Nichrome wires (17.78 µm diameter, A-M Systems, USA), which were gold-plated to reduce the impedance to 250 – 350 kOm (controlled by nanoZ, Plexon Inc., USA). Then, a commercially available Halo-5 microdrive (Neuralynx, USA) was assembled according to the producer instructions, where four single tetrodes were put into polymicro tubes (inner diameter 0.1 mm) and glued to the plastic shuttles. The microdrive was implanted and fixed first with quick adhesive silicone (Kwik-Sil, World Precision Instruments, USA) and then with dental cement (Henry Schein Krugg Srl, Italy).

After the surgery, the chicks were left to recover until the next day in their home cages. Between the 8th and the 12th day after hatching we recorded neural responses to face-like stimuli in the NCL of chicks. Before every recording session the microdrive was connected to the Plexon system (Plexon Inc., USA) via a QuickClip connector and an omnetics headstage (Neuralynx, USA). After every recording session the tetrodes were manually advanced by ca. 100 µm.

Signals were pre-amplified with a 16-channel head-stage (20×, Plexon Model number: PX.HST/16V-G20-LN) subsequently amplified 1000 × and digitalised. Spike detection and sorting was automatically performed in Kilosort 2.0^48^ with following parameters: ops.minfr_goodchannels = 0.1; ops.Th = [10 5]; ops.lam = 20; ops.AUCsplit = 0.95; ops.ThPre = 8; ops.spkTh = -6. All identified units were manually curated using Phy 2.0.

### Stimuli

To identify face-selective neural responses we used modified stimuli from previous behavioural experiments^4–5^ that showed innate preference of newborn chicks to face-like configurations (Fig. 1). An empty head silhouette (13 x 9 cm (H x W), 23.7° x 16.5° visual angle, Fig. 1A) was used for habituation and as a background image during recordings. Therefore, in every trial only the configuration of dots (1.1 cm, 2° visual angle) inside this head region (Fig. 1B-H, K-L) or a range of frequency components (Fig. 1I-J) of the stimulus was changed. We defined a proper face detector based on its stronger neural response to the face-like configuration (Fig. 1B) than to other stimuli with three dots (an “inverted” face and a “linear” configuration; Fig. 1C-D) or to single facial features (“eyes”, a “right eye”, a “left eye”, and a “beak”; Fig. 1E-H).

In addition, we tested how face neurons responded to asymmetrical “Picasso-like” images (Fig. 1K-L) with eyes shifted to the left/right to the medial sagittal line of the “upright” face-like stimulus. We also tested the response of potential face neurons to face-like stimuli dominated by high-frequency (>= 15Hz, Fig. 1I) or low-frequency (<= 5Hz, Fig. 1J) components, which corresponds to >= 15.05 cycles/face (>= 0.9 cycles/degree) and <= 4.95 cycles/face (<= 0.3 cycles/degree), respectively. To modify the frequency components of the face-like stimulus we used the “hard_filter” function in Matlab applied to the original “upright” face-like image of 1536 x 1063 pixels. The filter was applied separately to three colour channels, which were subsequently concatenated again.

During recording sessions, we randomly presented stimuli for 500 ms with inter-stimulus intervals randomly varying between 2500 and 3500 ms. Experiments were video-recorded using CineLAB system (Plexon Inc., USA). To enhance the motivation of birds to pay attention to the screen, random trials were occasionally rewarded (30-40%) by opening the feeder 500 ms after the stimulus offset for 500-1000 ms.

### Histological analysis

After the last neural recording birds were overdosed with the ketamine/xylazine solution (1:1 ketamine 10 mg/ml + xylazine 2 mg/ml). Electrolytic lesions were made at the recording sites by applying a high-voltage current to the tetrodes for 10-15 seconds. Then, the birds were perfused intracardially with the phosphate buffer (PBS; 0.1 mol, pH = 7.4, 0.9% sodium chloride, 5 °C) followed by 4% paraformaldehyde (PFA). Brains were incubated for at least two days in PFA and a further two days in 30% sucrose solution in PFA. Coronal 60 μm brain sections were cut at −20 °C using a cryostat (Leica CM1850 UV), mounted on glass slides, stained with the Giemsa dye (MG500, Sigma-Aldrich, St. Louis, USA), and cover slipped with Eukitt (FLUKA). Brain sections were examined under the stereomicroscope (Stemi 508, Carl Zeiss, Oberkochen, Germany) to estimate the anatomical position of recording sites.

## Data analysis

Based on the visual analysis of video recordings, we selected only trials in which birds looked at the stimulus with both eyes or with the eye contralateral to the recording site. For the final analysis, we considered only those units that were recorded for at least 10 trials for each stimulus type (32 ± 7 (mean ± standard deviation) trials per stimulus type).

### Identification of face-selective neural responses

The neural activity of recorded units was analysed in the 900 ms window starting from 100ms after the stimulus onset (to account for the visual latency of NCL neurons^49^) until 500 ms after the stimulus offset. First, every trial was smoothed using Gaussian kernel with 100 ms sigma. To identify stimulus-selective neural responses we then performed the sliding-window analysis of variance (one-way ANOVA; 10 ms bin window, 10 ms step-size) with the stimulus type as a factor (“upright”, “inverted”, “linear”, “eyes”, “left eye”, “right eye”, “beak”). For all significant (p < 0.01) time bins we performed a post-hoc analysis (“multcompare” function with Tukey–Kramer test) and selected only bins where the response to the tested stimulus was significantly different (p < 0.01) from other stimuli. If there was a significant response for at least a 100 ms period (10 consecutive bins) we performed an additional cluster permutation test to control for multiple comparisons. For this, all F-values within a significant response window were summed up (F-real) and compared to the sum of F-values resulting from the ANOVA and the post-hoc analysis of randomly shuffled trials (F-shuffled). This procedure was repeated 1000 times for every unit, and the response window was considered truly significant (i.e., face-selective) only if the F-real was higher than 95% of all F-shuffled (corresponding to the p < 0.05).

To estimate the probability of encountering false-positive responses in our dataset, we randomly sampled 224 trials (32 trials per 7 stimuli) out of all recorded trials (N=119567) for 1000 times and performed a sliding-window ANOVA with the post-hoc test for each shuffled dataset. We then compared the proportion of false-positive and real stimulus-selective units with the proportion test.

### Analysis of response properties of face-selective cells

To quantify the selectivity of individual face-responsive cells we calculated the face-selectivity index (FSI^50^) defined as the difference between the mean response to the face-like stimulus and the mean response to nonface stimuli, divided by the sum of these means. The mean response was defined as the average firing rate within the face-selective response window. FSI >= 0.33 or <=-0.33 corresponds to a 2:1 ratio and are considered to signify a strong face-to-nonface neural response.

We tested the sensitivity of identified face-selective neurons to violation of the symmetry of the face configuration. For this we compared the firing rate within the face-selective window between the “upright” face-like configuration and “Picasso-like” stimuli. We expected face-selective cells with inhibitory response (with the negative FSI) to show less inhibition in response to “Picasso-like” stimuli, while cells with the positive FSI to decrease their response to “Picasso-like” stimuli. Therefore, we used a Linear Mixed Effect model (LME) with “stimulus type” (“upright”, “picassoL”, “picassoR”) and FSI (“negative” or “positive”) as fixed effects and “unitID” (43 face-selective units) as a random effect. For the post-hoc analysis, we used the general linear hypothesis test (“glht” function from the “multcomp” package in R).

To investigate the response of face-selective units to different frequency components of the “upright” face-like stimulus, we compared the average firing rate within the face-selective response window between the high-frequency and the low-frequency stimuli. We did not perform a direct comparison between the frequency-modified and the “upright” stimuli, since luminosity and contrast between these stimuli differed and, thus, could be a confounding variable affecting the analysis. Therefore, we used the LME with “stimulus type” (“HF”, “LF”) and FSI (“negative” or “positive”) as fixed effects and “unitID” as a random effect. Pairwise comparisons were performed post-hoc using the general linear hypothesis test.

### Decoding of neural population responses

To quantify the amount of information about the face-like stimuli contained in the population response during the trial, we performed the percent-explained variance (PEV) analysis. PEV measures the percentage (ω²) of the variance explained by the tested factor. ω² is calculated from the sum of squares of the effect (SSeffect) and the mean squares of the within-group (error) variance (MSerror) (Equation 1).

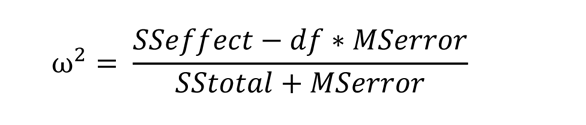

Based on coefficients from the sliding-window ANOVA performed for each stimulus-responsive unit we calculated ω² for each 10 ms bin of the analysed time window. To illustrate the explained variance for the entire population of units selective for a specific stimulus type, ω² values were averaged and compared to the average ω² obtained from the shuffled data (1000 times for each unit). We identified the populations’ information content to be significant when the ω² of real data was above the 95th percentile of the ω² from shuffled data (corresponding to p < 0.05).

To further evaluate how the neural population encodes face-like stimuli, we trained a multi-class support vector machine (SVM) classifier to discriminate between face-like and non-face stimuli based on the neural response of face-selective NCL neurons. We included neurons that selectively responded to the face-like stimulus and had at least 25 recorded trials per stimulus (N = 33 neurons). Every trial was smoothed using Gaussian kernel with 100 ms sigma. Then, we randomly selected 25 trials per stimulus type and divided them into the training set (23 trials per stimulus type) and the test set (2 trials per stimulus type). To make the neural activity comparable between units, it was subsequently z-scored using the mean and the standard deviation of the training set only. For a more robust estimation of the decoding accuracy, we performed one thousand iterations of the SVM training and testing, randomly selecting the trials each time.

We trained the multi-class linear SVM in Matlab with the following parameters: “one-vs-one” classification, 10-fold cross-validation, “linear” kernel function, “auto” kernel scale, with no additional regularization. We aimed, first, to test how similar are neural representations between different facial features. For this, we trained the SVM on the average neural responses during the face-selective window for each unit. Moreover, we aimed to estimate how the decoding accuracy changes during the stimulus presentation. For this analysis we performed the training and testing of the SVM for the whole duration of the trial divided into 10 ms bins, starting from 500 ms before the stimulus onset till 500 ms after the stimulus offset (overall duration 1500 ms).

The whole process of training and testing of the SVM has been repeated with randomly shuffled trials to estimate the proportion of false-positive predictions and compare it to the performance of the SVM trained on real data. For the network trained on the face-selective window the accuracy of the SVM predictions has been evaluated by the proportion test (p < 0.05 corresponds to the accuracy of >= 22%). For the SVM trained on the consecutive time segments within the trial range, the significant threshold for the prediction accuracy has been estimated based on the 95% confidence interval calculated for proportional data as 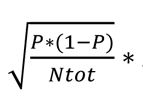 *Zscore*, where Pm is the proportion of correct predictions, Ntot is the total number of predictions, and Zscore is the inverse of the standard normal cumulative distribution function (“norminv” function in Matlab) evaluated at the probability values of 0.025 (for a two-tailed test).

All statistical analyses and visualization of the data was performed in R^51^ with packages “tidyverse”, “multcomp”, “ggplot2”, and “PMCMRplus” and in MATLAB using custom-made scripts.

## Acknowledgments

We are grateful to Anastasia Morandi-Raikova for her help with handling the chicks. This work was supported by funding from the European Research Council (ERC) under the European Union’s Horizon 2020 research and innovation program (Grant agreement no. 833504 SPANUMBRA) and from a PRIN 2017 ERC-SH4–A (2017PSRHPZ) to G. V.

## Author Contributions

D.K, O.R.S. and G.V. designed the experiment. D.K. performed the experiments. D.K. and M.Z analyzed the data. D.K. wrote the original draft of the manuscript and implemented the comments of all authors. All authors revised and edited the manuscript.

## Competing interests

Authors declare no competing interests.

## Materials & Correspondence

Correspondence and material requests should be addressed to G.V.

## Data availability

All data and code for analyses are available in the main text or in the depository (DOI 10.5281/zenodo.10517792).

## Notes

https://zenodo.org/records/10517793

### Competing Interest Statement

The authors have declared no competing interest.

