## Supplementary materials for "Innate face detectors in the nidopallium of young domestic chicks"

* Corresponding author: Giorgio Vallortigara

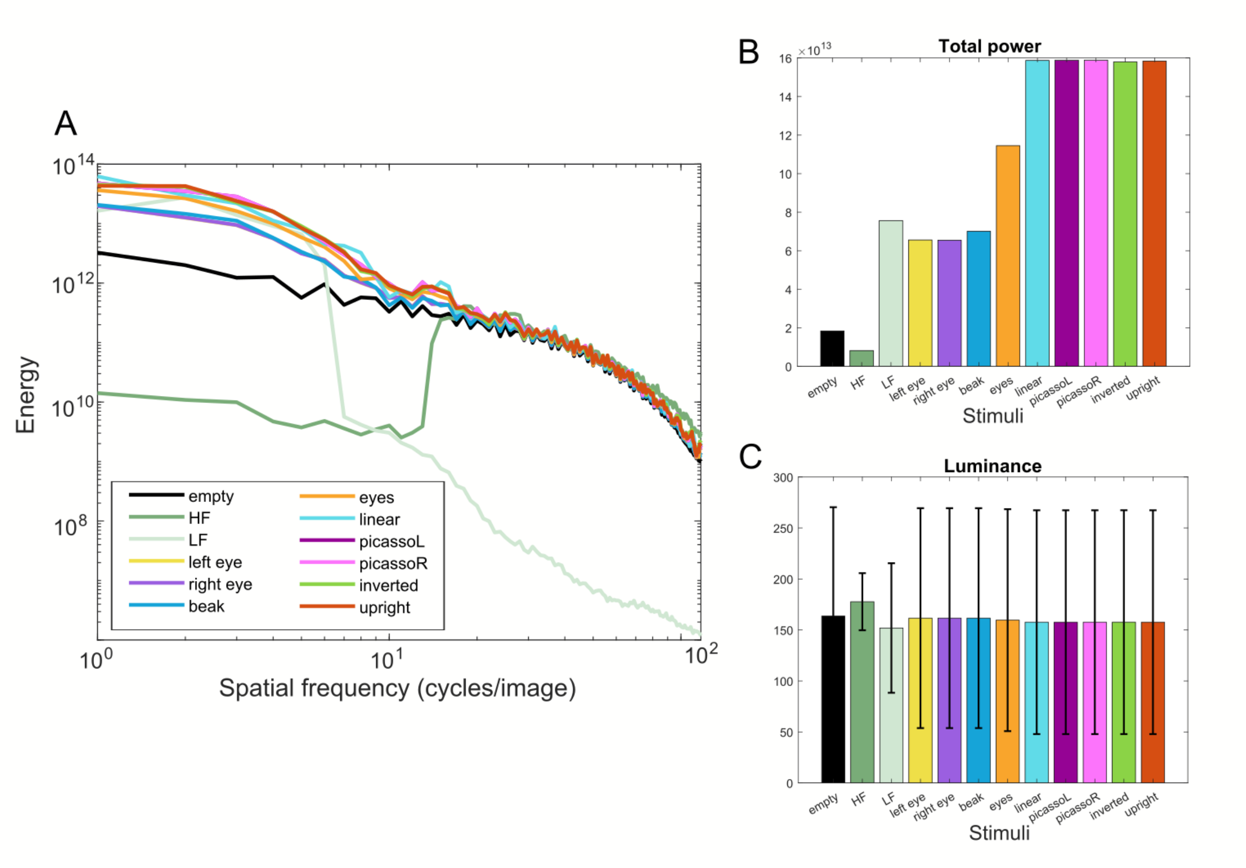


**fig. S1.**

**The schematic face-like stimuli have allowed us to better control for low-level visual features.** (**A**) The spatial frequency power spectra of all stimuli, except frequency-filtered "LF" and "HF", are very similar. The spectrum is shown up to 117 cycles/image (7 cycles/degree), which corresponds to the acuity limit of a domestic chicken[52]. (**B**) The total power of the frequency spectra shows that all stimuli featuring three dots ("inverted", "linear", "picassoL", "picassoR", "upright") are virtually identical, which indicates that the differences we observe in the neural responses to these stimuli cannot be explained by any difference in the frequency composition. (**C**) The average luminance of face-like stimuli is also very similar between different configurations. This can be explained by the fact that a difference in luminance can occur only from removing one or two black dots representing facial features (except in frequency-filtered stimuli). This modification can reduce the overall luminosity by 1 – 2% maximum since each dot covers only 1% of the overall stimulus area. The error bars correspond to the standard deviation of luminosity, which correlates with the contrast. Hence, the reduced contrast of “LF” and “HF” stimuli can account for some of the difference observed between the neural response to these and other face-like stimuli.

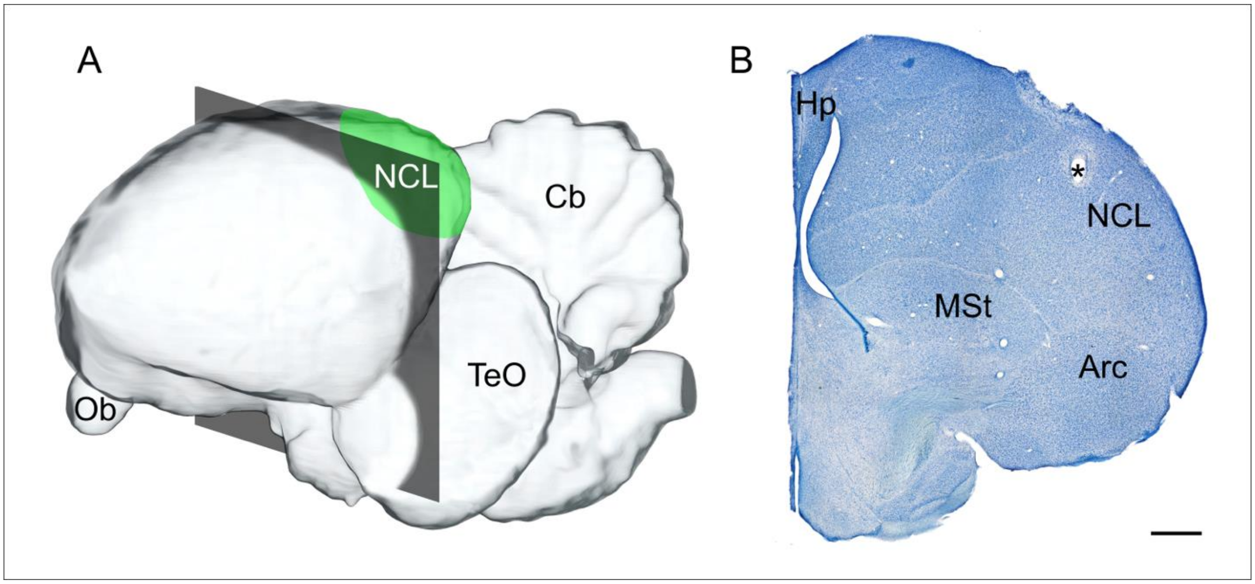


**fig. S2.**

An example of a recording site within the caudolateral nudopallium (NCL) of young domestic chicks. (A) A schematic representation of the avian brain, showing the approximate location of the NCL (in green) and the rostro-caudal position of the brain section presented on the right (grey-shaded area). (B) The Giemsa staining of the brain section shows the electrical lesion's location at the recording site (marked with an asterisk). Scale bar: 1mm. Abbreviations: Arc – arcopallium, Cb – cerebellum, Hp – hippocampus, MSt – medial striatum, NCL – caudolateral nidopallium, Ob – olfactory bulb, TeO – optic tectum.

**Movie S1.**

Neural activity of an exemplary face detector in the NCL of a young domestic chick shows selective response to the “upright” face configuration compared to other configurations and single facial features. The video recording of the trials corresponds to the neural activity (below) recorded from one electrode. The neural data are filtered between 300 and 6000Hz. The corresponding visual stimuli are shown in the right corner. The recording is slowed down to the half of the actual speed.
